# The acquisition of additional control over quorum sensing regulation buffers noise in microbial growth dynamics

**DOI:** 10.1101/2024.07.26.605310

**Authors:** Marco Fondi, Christopher Riccardi, Francesca Di Patti, Francesca Coscione, Alessio Mengoni, Elena Perrin

## Abstract

Quorum sensing (QS) is a cell-to-cell communication system used by bacteria to act collectively. Often, bacteria possess more than one QS regulatory module that form complex regulatory networks. Presumably, these configurations have evolved through the integration of novel transcription factors into the native regulatory systems. The selective advantages provided by these alternative configurations on QS-related phenotypes is poorly predictable only based on their underlying network structure. Here we show that the acquisition of extra regulatory modules of QS has important consequences on the overall regulation of microbial growth dynamics by significantly reducing the variability in the final size of the population in *Burkholderia*. We mapped the distribution of horizontally transferred QS modules in extant bacterial genomes, finding that these tend to add up to already-present modules in the majority of cases, 63.32%. We then selected a strain harboring two intertwined QS modules and,using mathematical modelling, we predicted an intrinsic ability of the newly acquired module to buffer noise in growth dynamics. We experimentally validated this prediction choosing one strain possessing both systems, deleting one of the two and measuring key growth parameters and QS synthase expression. We extended such considerations on two other strains naturally implementing the two versions of the QS regulation studied herein. Finally, using transcriptomics, we show that the de-regulation of metabolism likely plays a key role in differentiating the two configurations. Our results shed light on the role of additional control over QS regulation and illuminate on the possible phenotypes that may arise after HGT events.

## Introduction

Quorum sensing (QS) is a communication system among bacterial cells that allows a coordinate collective behavior depending on cell density, growth rate, and on the composition of the surrounding microbial community [1, 2, 3]. It is based i) on communication molecules of different chemical nature, called autoinducers (AI), that are produced and secreted by bacterial cells and ii) on receptor regulators that, after the recognition of a given amount of a specific AI, can control the expression of several different genes [4]. In Gram-negative bacteria QS is typically regulated by N-acyl-homoserine lactone (AHL) molecules-mediated systems. This is the case of the model organism *Vibrio fischeri* where LuxI synthesizes an AHL signal and LuxR is an AHL receptor protein that activates or represses gene expression by binding to a consensus sequence in the promoter regions of target genes [5]. Many different cellular processes are under the control of QS, including virulence, pathogenesis, biofilm formation and fluorescence [6]. They are generally highly expensive from an energetic point of view, and bacterial cells activate them only when a large population can benefit from them. As a consequence, the regulation of this phenotype is tightly controlled by microorganisms that, in time, have evolved many different regulatory strategies that ultimately give origin to complex behaviors that include environment-dependent life-style switches [7]. Interestingly, in many cases, bacteria possess more than one QS system [4, 6, 8].While in some cases the multiple QS systems are based on distinct types of signal molecules, in others they respond to the same class of compounds, thus giving rise to complex regulatory interactions whose outcomes still not understood. Arguably, these complex regulatory systems derive from simpler systems (e.g. a native LuxIR module) that were later modified by diverse evolutionary processes. Early studies on the origin and evolution of AHL-based systems, for example, have indicated HGT as a possible factor in the spreading of QS modules among Bacteria [9, 10]. Nowadays, it is clear that QS genes are indeed often associated with mobile genetic elements (MGEs), that QS circuits can move through HGT [11, 12, 13] and that upon transmission they will begin interacting with the residing ones. Much less clear and predictable are the effects of these non-native QS modules on the overall phenotype of the receiving cells, despite they might be of importance in the characterisation of specific biological processes (e.g. infections, biodegradation, etc.) as shown by Bellieny-Rabelo et al. (2020). These authors showed that the HGT-mediated reprogramming of QS regulatory circuit, conferred specific selective advantages for the expression of crucial host adaptation- and fitness-oriented systems in the *Pectobacterium* and *Dickeya* [14]. Similarly, representatives of the *Pseudoalteromonas* genus were shown to possess highly heterogeneous QS regulation schemes with QS modules that resemble those of other microorganisms (like *Escherichia coli*, *Vibrio* and *Pseudomonas* species). The addition of these modules has been hypothetically linked to the capability of colonizing complex ecological niches thanks to a more finely tuned, QS-mediated, collective behavior [15]. From a broader perspective, the inclusion of a regulatory module inside a mobile element and its further acquisition by a microorganism, represents an extraordinary example of the plasticity of cellular transcriptional networks and an exceptional occasion to study the effect of alternative regulatory architectures on the expression of some phenotypic trait(s). Indeed, as HGT and/or recombination events lead to the appearance of novel regulatory modules, these, to be maintained over evolutionary time scales, have i) to integrate with the native one(s) and, eventually, ii) increase the fitness of the recipient organism. As for the first point, as stated, recent analyses confirmed that some extant QS regulatory networks engaged in HGT during their evolution and have been maintained by present-day microorganisms. Regarding the second aspect, instead, our knowledge is lagging far behind. In other words, it remains unknown why the regulation of QS in some microorganisms is hierarchical (i.e. composed by multiple, interacting modules as shown, for example, in the genus *Burkholderia* [16, 17] or in *Vibrio fischeri* [18]) and which are the functional advantages of multi-input (QS) systems. Here, we first quantified the distribution of non-native LuxI and LuxR copies across Prokaryotes, validating the role of HGT in shaping the structure and dissemination of QS modules within microbial genomes. To gain a deeper understanding of the impact of acquiring an additional QS module on the phenotype of a receiving cell, we then selected one specific case among those identified in our genome-level analysis (the CepIR and CciIR systems in *Burkholderia*) and used mathematical modelling to predict the possible outcomes of introducing an extra QS module in a genome. To confirm the model predictions, we generated mutants harboring a reduced (*core*) regulatory circuit (mirroring the pre-HGT situation) and compared their QS response with the wild-type circuit (*complete* circuit, mirroring the post-HGT architecture). Finally, whole transcriptome sequencing and differential gene expression analysis identified key metabolic genes repression as a possible effector in diverting the phenotype in cells harboring the additional QS module. Overall, we observed the capability of the newly acquired module to buffer noise in cellular growth dynamics and, from a more general standpoint, we addressed the possible consequences of evolution-driven changes in the structure of a regulatory circuit.

## Results

### The occurrence of non-native QS-modules in prokaryotic genomes

We started by examining how often a QS module composed of a *luxIR*-like gene pair is found on genomic islands (GI) i.e. large chromosomal regions with evidence of HGT. This put our work into perspective since a large body of literature supports the idea that the members of the canonical QS module (i.e. a *luxIR* gene pair) share a common origin followed by co-evolution as regulatory modules [19]. Because many bacterial strains contain multiple QS modules in their genome, HGT from independent sources has been indicated as a possible way to expand/modify the underlying regulatory circuits [12, 9, 20, 19]. To verify this hypothesis and quantify the contribution of HGT in this process we searched present day GIs for the presence of QS modules. Overall, we identified 259 microbial GIs containing at least one LuxI-like and one LuxR-like coding genes within close boundaries (Figure **1**, **Supplementary File S1** and 10.5281/zenodo.12796961). Conversely, we found 3087 LuxIR-like pairs that were i) close to one another in their harboring genome and ii) were not included in any of the GIs analyzed. This latter group of sequences likely represents a set of native LuxIR pairs, i.e. QS modules that did not originate from a transfer event, at least in evolutionary recent times. In terms of relative frequency that’s 8% of 3,346 analyzed QS modules associated to a GI. Perhaps more importantly, in 63.32% of the cases a newly acquired QS module entered a chromosome that already owned a QS module, thus probably giving rise to a complex interplay between the native and the non-native regulatory systems. These results underscore the importance of HGT in determining the distribution of added QS modules in extant prokaryotic genomes. However, what remains to be addressed is the possible array of selective advantages provided by the acquisition of additional QS modules. In other words, why would multiple, horizontally transferred QS modules be maintained over the course of evolution? In the following paragraphs we will describe in detail one specific case of non-native QS modules integration among those shown in Figure **1** and examine its possible contribution to specific cellular phenotypes.

**FIG 1.**
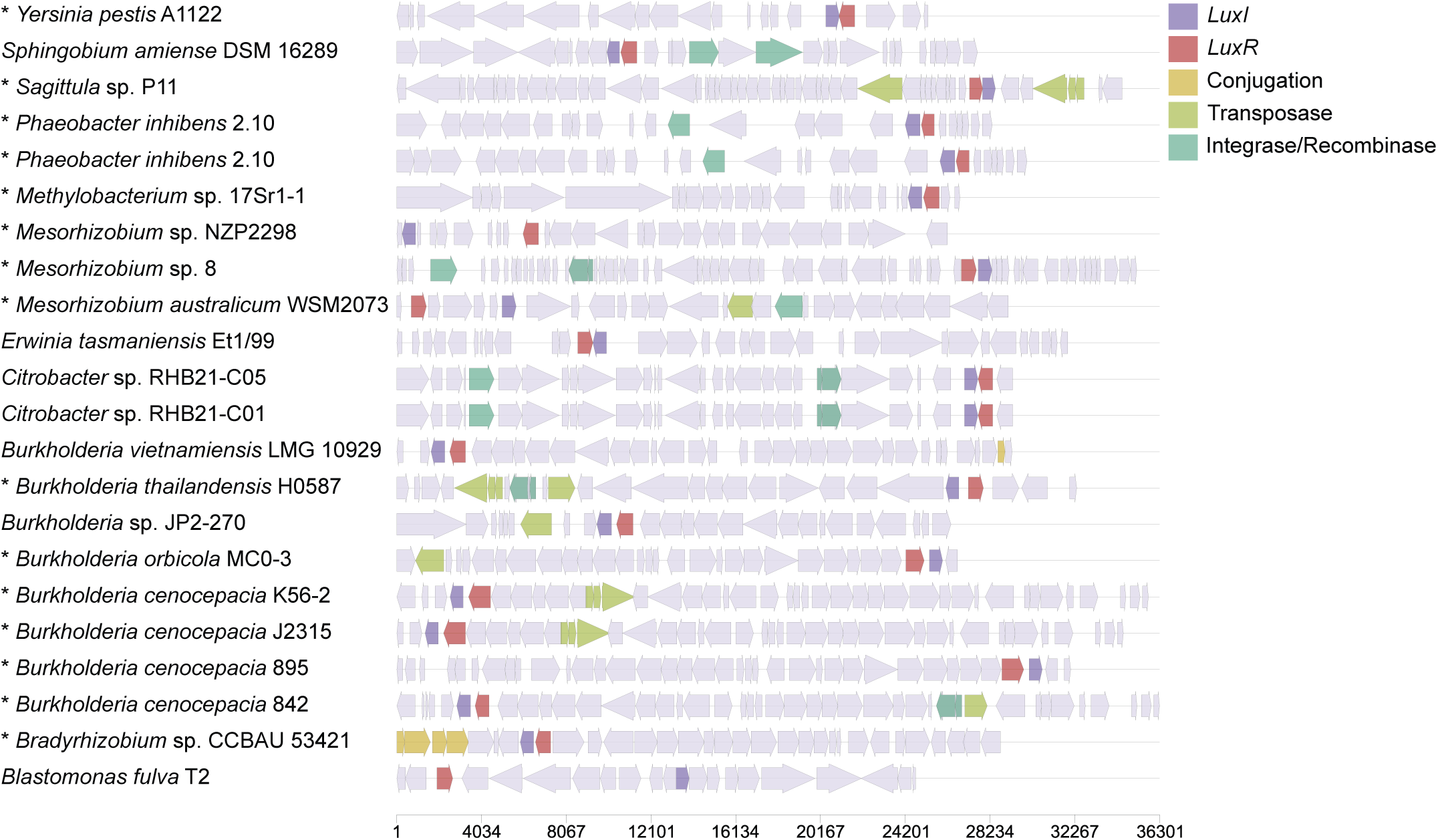
A sample of 22 *luxIR*-like containing genomic islands from our dataset. The direction of each arrow indicates whether that feature is annotated on the forward strand (pointing right) or reverse strand (pointing left). Marked by an asterisk are those that have at least another LuxIR couple elsewhere on the native genome. The fourth and fifth GIs from the top are the same species yet come from two distinct assemblies (GCF_003443555.1 and GCF_000154745.2, respectively). LuxIs are shown in purple and LuxRs in red, whereas protein-coding genes involved in genetic mobilization are displayed in shades of orange and green. The representation of the entire set of 259 GIs identified in this study are available as **Supplementary File S1**.

### Two concurrent QS regulations in the *Burkholderia* genus

Among the QS modules with a history of HGT identified in the first part of the analysis, we picked as study case the well characterized CepIR and CciIR systems in *Burkholderia* [21]. Besides the standard LuxIR-like system that goes under the name of CepIR, some of the strains in this genus possess an integrated regulatory module named CciIR that influences (and is influenced by) CepIR expression, as schematically shown in Figure **2**A. Specifically, the integration of CciIR and CepIR systems occurs through the implementation of several positive and negative control loops (Figure **2**B). The overall layout of the CepIR and CciIR QS modules resembles that of the classical LuxIR circuit, except for the negative autoregulation of CepR and CciR (LuxR-like proteins are commonly found as positively autoregulated). Furthermore, being associated to a genomic island (*cci* island), this additional module is prone to being horizontally transferred, giving rise to a potentially patchy genomic distribution and, in turn, to a *complete* (CepIR and CciIR systems together) vs. *core* (CepIR only) organization of QS regulation in this group of microorganisms. We assessed the distribution of the *complete* and *core* architecture across all reference species of *Burkholderia* by mapping in silico the presence/absence patterns of the *cci* genomic island genes. Despite part of the *cci* genomic island being used as an epidemic strain marker [22], its actual distribution inside *Burkholderia* has not been investigated at the genus level. For this reason, we probed the presence of the *cci* island encoded genes (55) in all the available complete *Burkholderia* genomes and combined this information with the genomic relatedness of the representatives of this genus. We measured the relatedness in terms of Average Nucleotide Identity (ANI, see Methods) for each pair of *Burkholderia* genomes included in the dataset. As reported in **Supplementary File S2**, the distribution of the genes by the *cci* island follows a pattern that matches the one obtained through ANI computation. More in detail, in the cluster comprising *B. mallei*, *B. pseudomallei* and *B. thalilandiensis* species, the complete set of *cci* genes is never found. Instead, the distribution of *cci* genes in the other cluster (hereinafter *cci*-group, that includes, among the others, the *Burkholderia cepacia* complex (Bcc) and the members of the *B. glumae/gladioli/plantarii* group) is patchy and includes (22) microbes that harbor more than 50% of the reference *cci* genes and others (93) possessing less than 50% of the reference *cci* genes (Figure **2**C). This pattern suggests a complex evolutionary history for this genomic island, mainly guided by gain/loss events that resulted in some strains harboring a *complete* architecture (e.g. *Burkholderia cenocepacia* J2315, K56, ST32) and others a *core* one (e.g. *B. cenocepacia* AU1054, H111, etc.), as schematically shown in Figure **2**C. Having defined the taxonomic boundaries of the QS modules distribution in *Burkholderia*, we next asked which may have been the consequences and advantages (if any) provided by the acquisition of a novel, additional QS control system and its integration into the native regulatory circuit.

**FIG 2.**
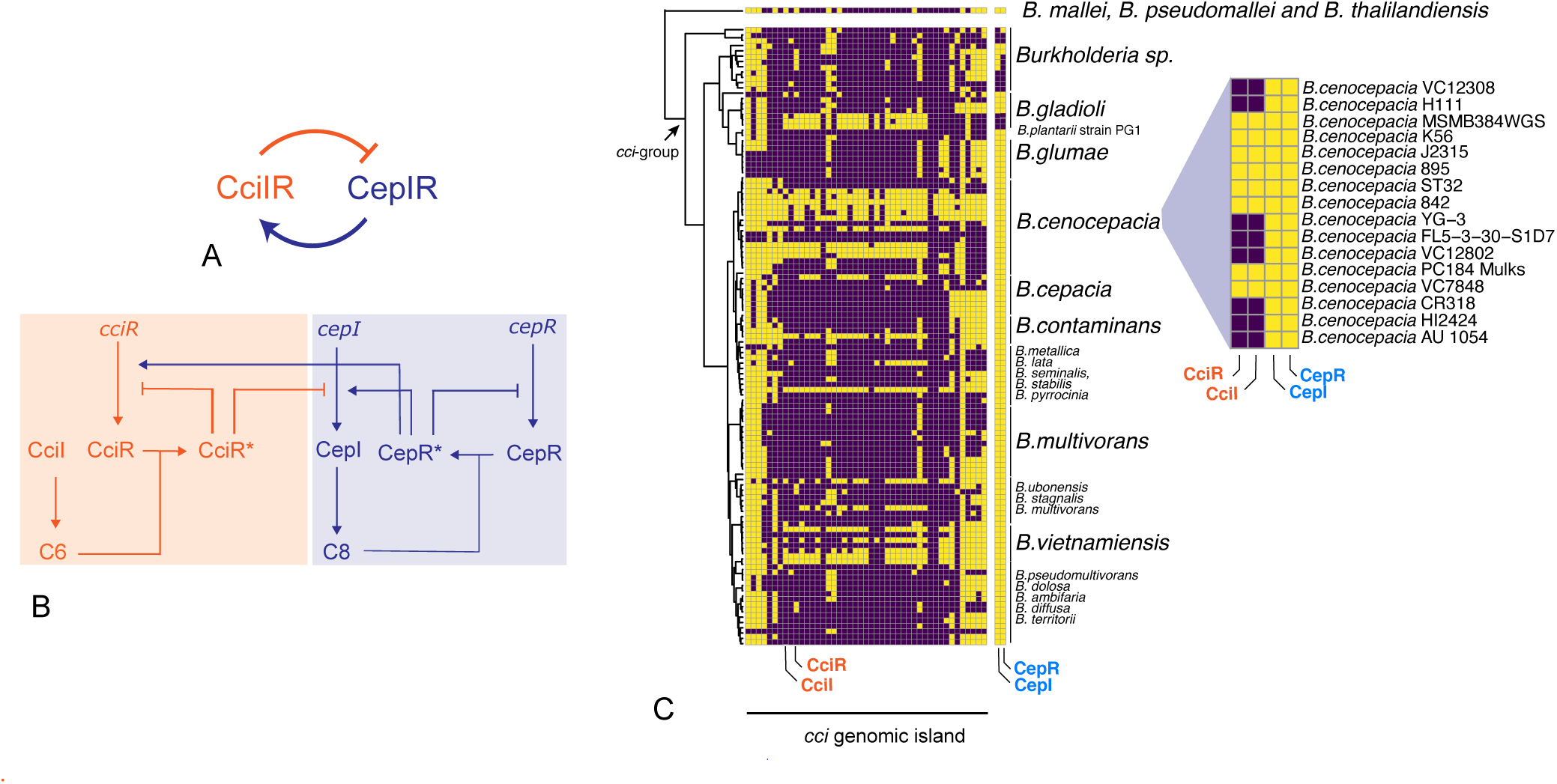
A) Schematic representation of the relationship betweeen CepIR and CciIR QS modules. B) Detailed representation of the interconnections between CepIR and CciIR QS modules. C) Phylogenetic distribution of the *cci* genes in the *Burkholderia* genus with a focus on the CepIR and CciIR distribution in *B. cenocepacia* species. Yellow and purple sqares indicate the presence and the absence of the corresponding genes, respectively.

### Modelling the effects of *core* and *complete* architectures

To address this point, we implemented a double strategy. First, we selected one of the strains that naturally bears a *complete* configuration (*B. cenocepacia* K56-2) and removed its *cciIR* module. Gene deletion was confirmed using whole-genome sequencing as detailed in **Supplementary File S3** and shown in **Figure S1**. We monitored key physiological features including growth rate, *cepI* expression dynamics and overall cell growth over a time period of 48 hours for i) a wt *B. cenocepacia* K56-2 strain harboring both QS modules (*complete*) and ii) a mutant strain bearing the CepIR QS module only (*core* architecture). Using a Lux reporter system (see Methods), we first monitored the expression of the AHL synthase coding gene *cepI* in the two strains. The dynamics of *cepI* expression was quite similar in the two strains (Figure **3**A). Both strains showed similar *cepI* expression values at the beginning of the experiment but, while in the wt *cepI* expression stopped around hour 15, the cells harboring the *core* architecture stopped expressing *cepI* about five hours later. Thus, the *core* architecture led to a higher maximal *cepI* expression with respect to the *complete* (wt) configuration, as shown in Figure **3**C. The observed difference was statistically significant (Wilcoxon rank sum test, p-value *<* 2.2e-16). Following this expression peak, both strains stopped expressing *cepI* and promoter activity was not detectable around hour 48. Concerning the growth dynamics of the two strains, we did not observe any difference in their maximal growth rates, and in the range 0 to 30h the two growth curves exhibit a similar trend, Figure **3**B). Afterwards, we formalized the two QS regulatory networks (*core* vs. *complete*) into a comprehensive mathematical model. The QS model implemented in this work is schematically represented in Figure **2**B and detailed in **Supplementary File S3**. The model takes into account the main players shown in Figure **2**B, i.e. 4 genes (*cepI, cepR, cciI* and *cciR*), 4 proteins (CepI, CepR, CciI and CciR), 2 activated transcription factors (CepR* and CciR*) and 3 metabolites (C8HSL, C6HSL, O-ACP, octanoyl acyl carrier protein). The latter represents the most likely substrate for the production of AI molecules [23]. The entire model (hereinafter termed *complete*) recapitulates the architecture of the *Burkholderia* representatives possessing both the CepIR and CciIR modules (Figure **2**C), whereas the architecture of those species harboring only the CepIR regulation is accounted for by the *core* model (the blue circuit of Figure **2**B). The *complete* model contains 18 parameters (**Supplementary File S3, Table S1 and S2**) and 11 of them are shared with the model accounting for the *core* architecture, thus making the *core* a subset of the *complete* model. Growth and *cepI*/*cciI* expression were linked through a Hill-like kinetic in which the growth of the bacterial cells is gradually stopped by the increase of the autoinducer concentration produced by the population (see **Supplementary File S3**). We then used the experimental data (namely *cepI* promoter activity, a proxy for *cepI* expression) and OD (a proxy for cellular growth) from the mutant strain to derive the most likely values for the model parameters using a stochastic curve-fitting method, as described in the Methods section (Figure **3**C). The set of the fitted parameters of the *core* model is reported in Table S1. We then proceeded to fit the *complete* model. Initially, we constrained the parameters of the *complete* model shared with the *core* model to their fitted values. However, this produced an unsatisfactory fit (dashed line in Figure **3**C) and, for this reason, we decided to remove the constraints of the values obtained when fitting the *core* model. By doing so, the quality of the fit improved and the models accurately recapitulated the overall dynamics of the system (Figure **3**C and D). More specifically, we computed the goodness-of-fit in both cases by resampling 1000 times the array of the predicted values (to have arrays of simulated and experimental data of the same length) and calculating the coefficient of correlation between the experimental data and the prediction for each of the resamplings. When considering all the resamplings, we found a better (and statistically significant) fit for the model with unconstrained parameters (0.90 vs 0.88, p-value *<* 0.0001, t-Test). This is compatible with a deep rewiring of the QS regulatory circuit following the deletion of one of the two modules in *B. cenocepacia* K56-2, possibly suggesting a deep and strong integration and feedback between the two systems. The values of the parameters together with their confidence intervals for both the *complete* and *core* models are reported in **Supplementary File S3, Table S1 and S2**.

**FIG 3.**
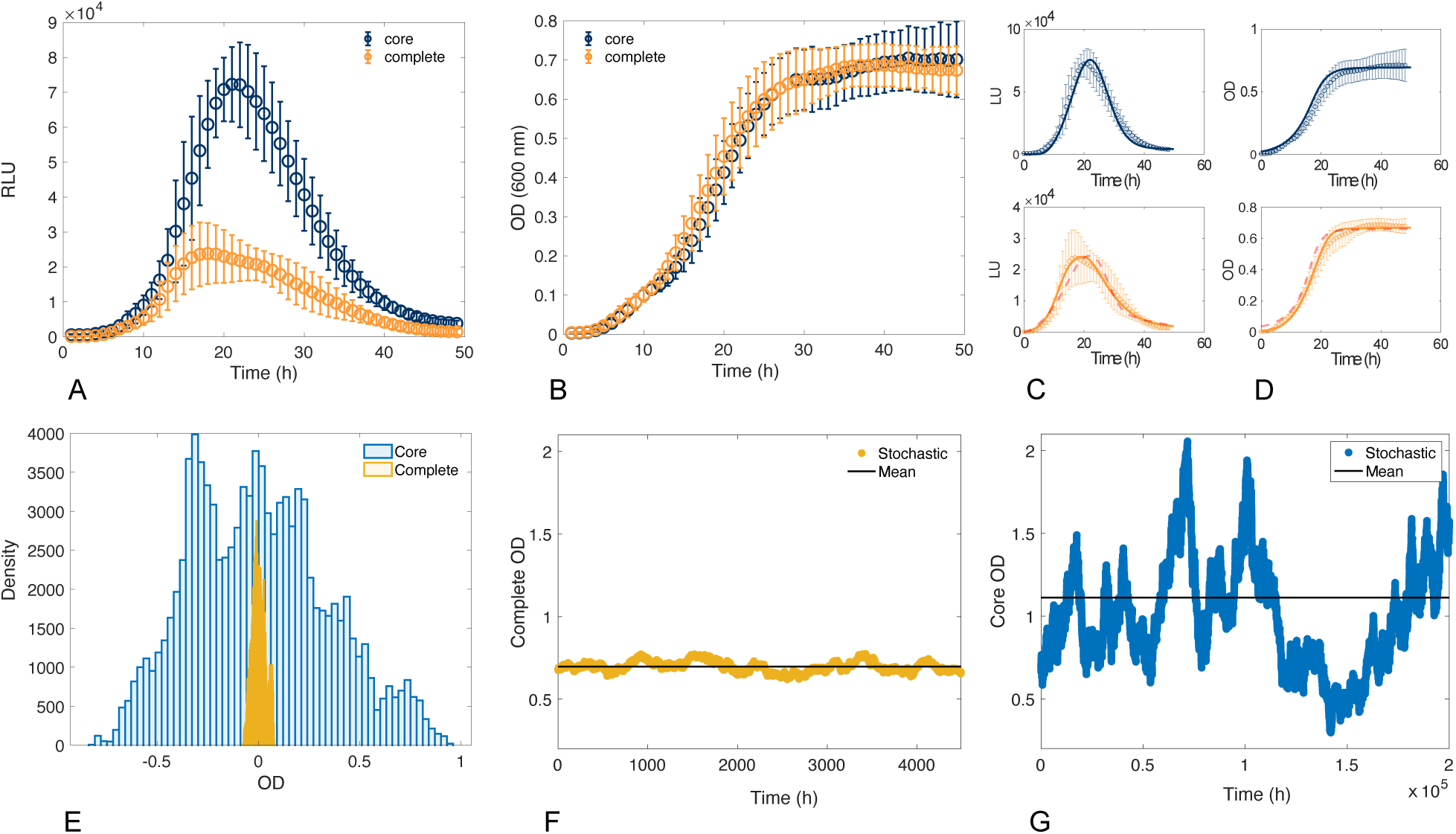
A) *cepI* and *cciI* expression in the *core* and the *complete* architectures. B) OD values in the *core* and *complete* architectures C) Model fit of the *complete* and *core* models for *cepI*. D) Model fit of the *complete* and *core* models for OD. In C9 and D), experimental data are represented by empty circles and error bars while simulations are represented by solid lines. E) The comparison between the fluctuations about the steady state of OD in the *complete* and the *core* model. F) A single stochastic realization implemented through the Gillespie algorithm [24] for OD values in the *complete* model. G) A single stochastic realization implemented through the Gillespie algorithm for OD values in the *core* model.

While the deterministic approach is key to understand whether the model is capable of reproducing the main features of these regulatory circuits, it cannot take into account one of the key characteristics of QS, i.e. the capability of cells to sense the correct signal concentration and behave in a coordinate fashion, in the presence of fluctuations. These are known to deeply affect cell regulation and signalling [25, 26, 27]. Stochasticity in biological circuits plays a pivotal role in determining the inherent variability among individual cells within a population. Cell-to-cell heterogeneity arises from intrinsic stochastic events in gene expression, protein synthesis, and cellular processes. This variability impacts diverse phenomena, from developmental processes to disease progression and cells have been shown to be able to exploit/control different sources of heterogeneity, depending on the situation [28, 29]. We sought to investigate whether the two architectures could had any role in controlling noise and/or perform noise buffering on cellular growth. To capture this inherent randomness and account for the diverse behaviors observed at the population level in our experiments, we simulated the stochastic model described by equations (1)-(26) in **Supplementary File S3** using a stochastic algorithm [24], specifically focusing our attention on cell density at stationary level. We computed the fluctuations of the predicted OD values in respect to the steady state level in the *core* vs. the *complete* models (Figure **3**E-G). The results of the stochastic simulations revealed that fluctuations about the steady state are much more pronounced in the *core* model, while the stochastic trajectories of the *complete* model span over a much narrower range (Figure **3** E-G). Overall, the model seems to suggest that the steady-state concentration of the species in the *complete* model is less affected by stochastic fluctuations. On the contrary, the *core* model appears to be more sensitive to random changes in the concentration of the different species of the model, reflected by larger fluctuations around their stationary values.

### An additional feedback loop over QS regulation influences population-level growth heterogeneity

To validate the model predictions and to detect noise, if any, in the final stages of *Burkholderia* growth we repeated the experiment described before (shown in Figure **3**A and B) for a total number of 27 replicates (Figure **4**A-D). Increasing the number of measurements and evaluating the final OD at the end of each growth period reached by cells harboring the *complete* vs *core* architecture we found a large, statistically significant variability in the latter compared to the former (Levene’s test of variances, p*<*0.0025, Figure **4**D). We also tested whether the difference between the sample variances resulted to be significant across those time points (i.e. after 35 hours). In that case the OD values followed normal distributions (p-value = 0.8732 and 0.1409 for *complete* and *core* according to Shapiro-Wilk normality test, respectively) and their variances were tested with an F-test (p-value = 1.438e-05). Prompted by this finding, we also tested whether, besides cell density, also *cepI* expression displayed heterogeneous expression values among the replicates. No statistically significant differences were found for the variance of *cepI* expression between mutant and wt strains **4**A and B). From these results we conclude that one of the differences between the strains harboring the *core* and the *complete* architectures resides in the capability of the latter to efficiently adjust the number of cells around a well-defined value. *Core* circuit-harboring cells lack this fine tuning, thus showing a much more variable signal in terms of cell growth (Figure **4**D). As QS is generally considered to be a cell density dependent process, we tested whether the capability of the *complete* circuit to stabilize the final cell density on a certain value was dependent on the initial number of cells of the population (similar to an *inoculum effect*). We evaluated, over three independent experiments, the growth dynamics and the *cepI* promoter activity from a range of initial values of cell density. For the sake of clarity, here we present normalized OD data in which each OD value has been divided by the maximum OD value observed in the corresponding experiment (raw data from this experiment are reported in **Supplementary File S3, Figure S2**). As shown in Figure **4**E, the final cell density of cell populations harboring the *complete* circuit was not influenced by the initial OD of the cultures. On the contrary, the mutant strains harboring the *core* configuration reached a much broader range of final OD values Figure **4**F. The robustness of this observation was successfully evaluated by testing the variance of the final endpoints of the two alternative configurations (*core vs. complete*, Levene’s test p-value = 0.0001414, Figure **4**G). Such narrower range of final cell densities for the *complete* vs. the *core* architectures was also observed when endpoints from the same starting OD were compared between the two configurations (**Supplementary file S3, Figure S3**). From these results, we conclude that, the acquisition of an extra QS module mitigates the variability of the final population cell density, independently from the initial number of cells within the tested range, at least in the strains and in the conditions tested herein. Finally, we checked whether the phenotype of strains naturally possessing one of the two regulatory schemes discussed before resembled those of the *B. cenocepacia* K56-2 wt and mutant strains. To this purpose, we randomly selected two more strains from the *B. cenocepacia* group shown in the inlet of Figure **2**C and accounting for the *core* (*B. cenocepacia* AU1054) and *complete* (*B. cenocepacia* J2315) structures. We then tested the variability of their end-point cell density, in relation to the initial number of cells (i.e. repeating the experiment whose results for the *B. cenocepacia* K56-2 wt and mutant strains are shown in Figure **4**E, F and G). As shown in Figure **5**, *B. cenocepacia* AU1054 displayed a significantly higher variability in the end-point OD600 in respect to *B. cenocepacia* J2315 (p-value *<* 0.00001, Levene’s test), a result that is in line with that obtained with *B. cenocepacia* K56-2 wt and mutant strains and the predictions of the mathematical model. A narrower range of final cell densities for the AU1054 vs. J2315 architectures was also overall observed when endpoints from the same starting OD were compared between the two configurations (**Supplementary file S3, Figure S4**).

**FIG 4.**
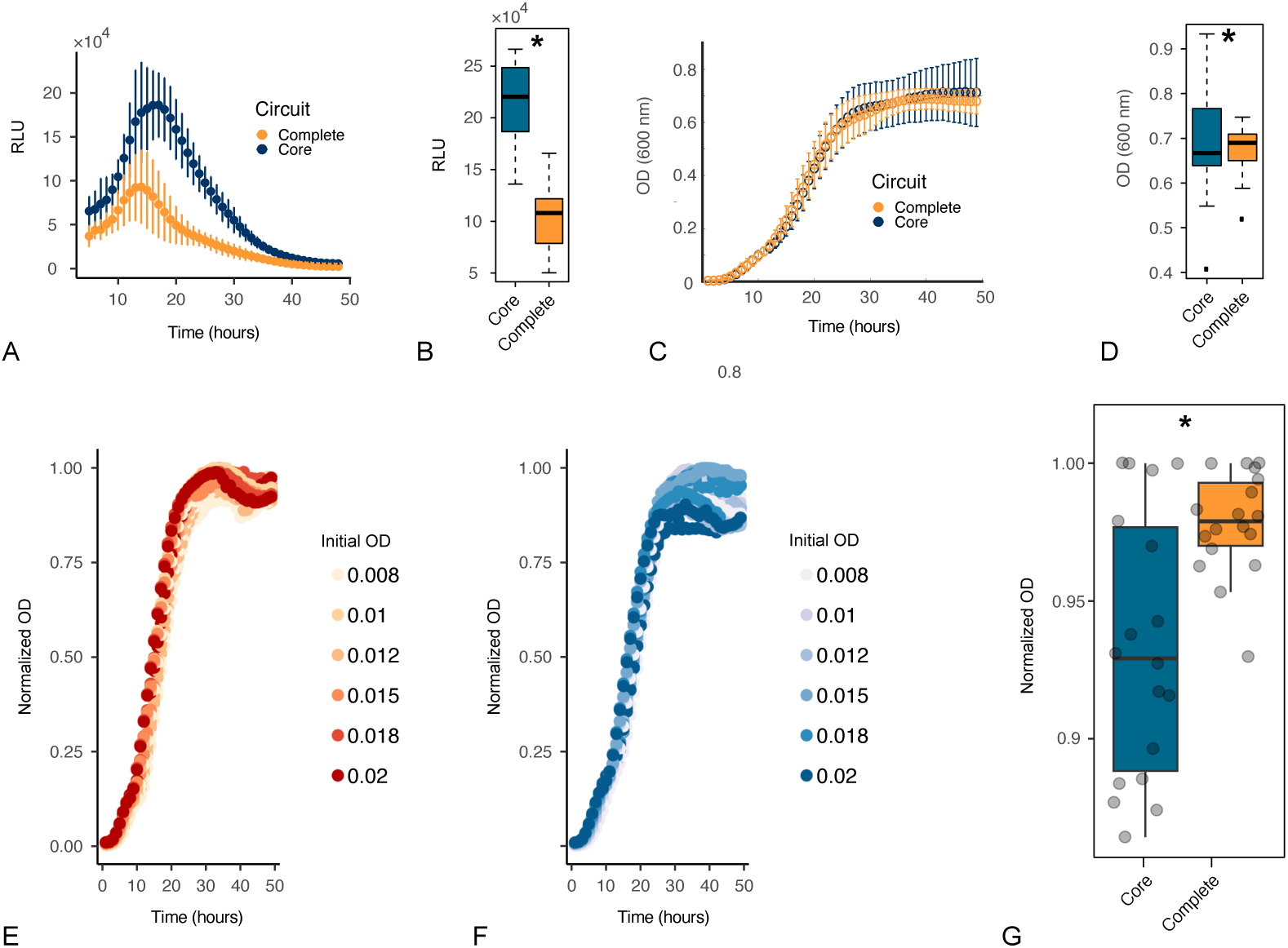
A) *cepI* expression in the *complete* and *core* configurations. B) Boxplot showing the variation across the maximal *cepI* expression values in the *complete* vs. the *core* configuration (the asterisk indicates a statistically significant comparison) C) Growth dynamics of strains harboring the *complete* vs. the *core* QS regulatory configurations. D) Boxplot showing the variation across the final (end-point) OD values in the strains harboring the *complete* vs. the *core* configuration (the asterisk indicates a statistically significant comparison). E) Growth curves for different starting OD values for strains harboring the complete and (F) the core configurations. G) Boxplot showing the variation across the final (end-point) OD values of plots in E and F panels (the asterisk indicates a statistically significant comparison).

**FIG 5.**
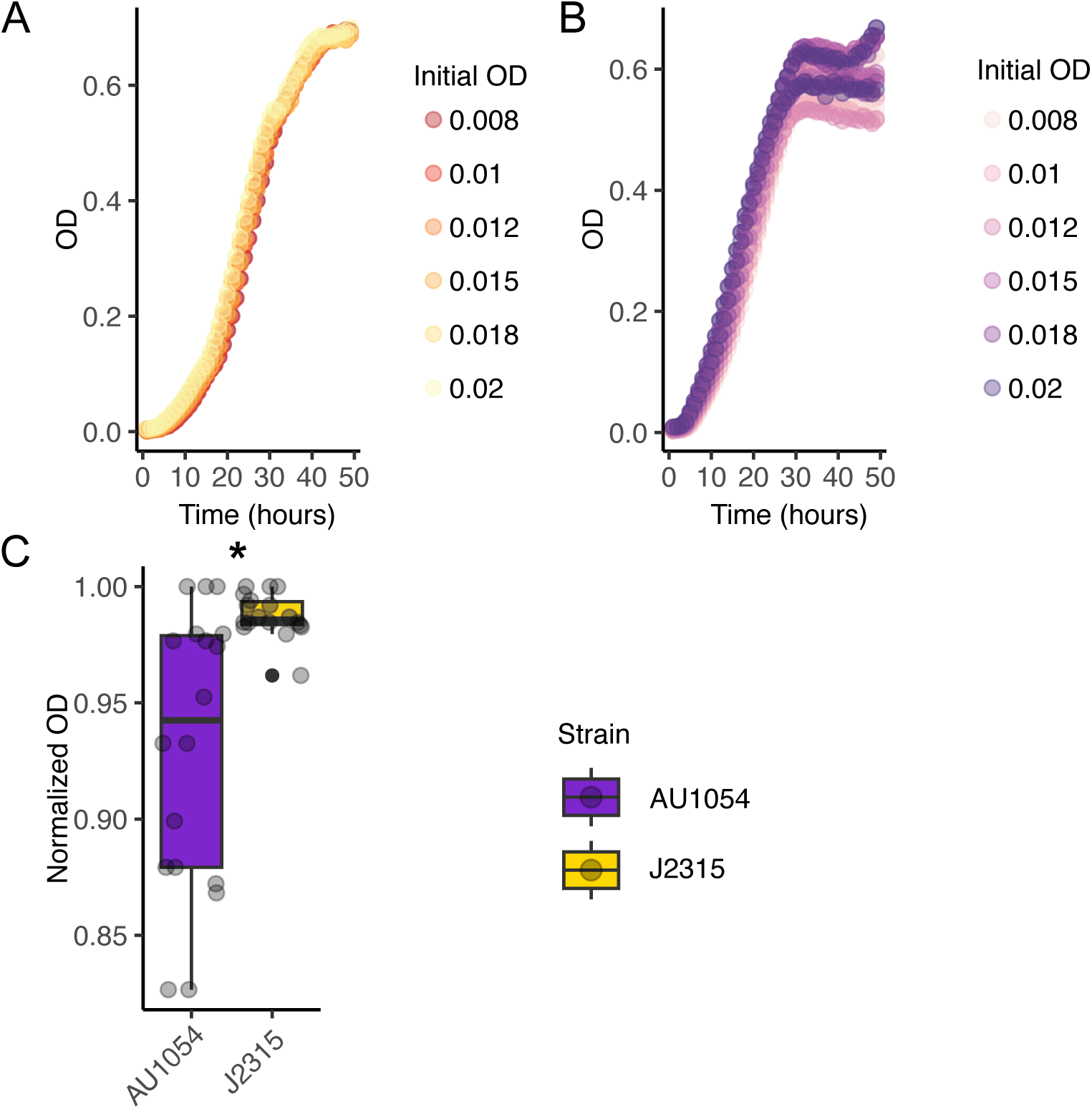
A) Growth curves for different starting OD values for *B. cenocepacia* AU1054 (*core*-like architecture). (B) Growth curves for different starting OD values for *B. cenocepacia* J2315 (*complete*-like architecture). C) Boxplot showing the variation across the final (end-point) OD values of plots in A and B panels (the asterisk indicates a statistically significant comparison)

### *Core* architecture leads to down-regulation of metabolism in the mutant strain

Finally, we investigated which cellular processes might influence the differing phenotypes observed between the *core* and *complete* strains (*B. cenocepacia* K56-2), and performed RNA-seq. We sampled the transcriptomes of the strains harboring the two architectures at three different time points of their growth curves, at 20, 25 and 30 hours. We focused on three time points closer to the end of the growth curves since that’s where most of the differences in terms of growth dynamics reside (Figure **4**C). The results of this experiment are shown in Figure **6**. After applying variance stabilization on the gene expression matrix we explored how well the different architecures over time explain patterns across all samples. Figure **6**A shows a clear distinction of the samples in a time-specific manner. Moreover, the deletion strain consistently localizes on top of / separated from the wild type. Some minor, local attractions are also observable. This is in line with the phenotypic differences among the strains that are particularly evident in the final stages of their growth curves (Figure **4**C). In all three cases the density is strong around a fold change of 0 (Figure **6**B, meaning that most of the genes are not differentially expressed or that they exhibit a mild change in gene expression). Samples at T3 show a more relaxed distribution with a slight offset towards negative log fold changes compared to time points T1 and T2 but similar decay around an absolute LFC of 1. We then filtered the genes by significance threshold (padj *<* 0.05, Wald significance test) and required that the absolute LFC be greater than 1. Table **1** summarizes the upregulated and downregulated genes in the mutant, at each time point, filtered by statistical significance (padj *<* 0.05, Wald significance test) and by absolute LFC be greater than 1.

**FIG 6.**
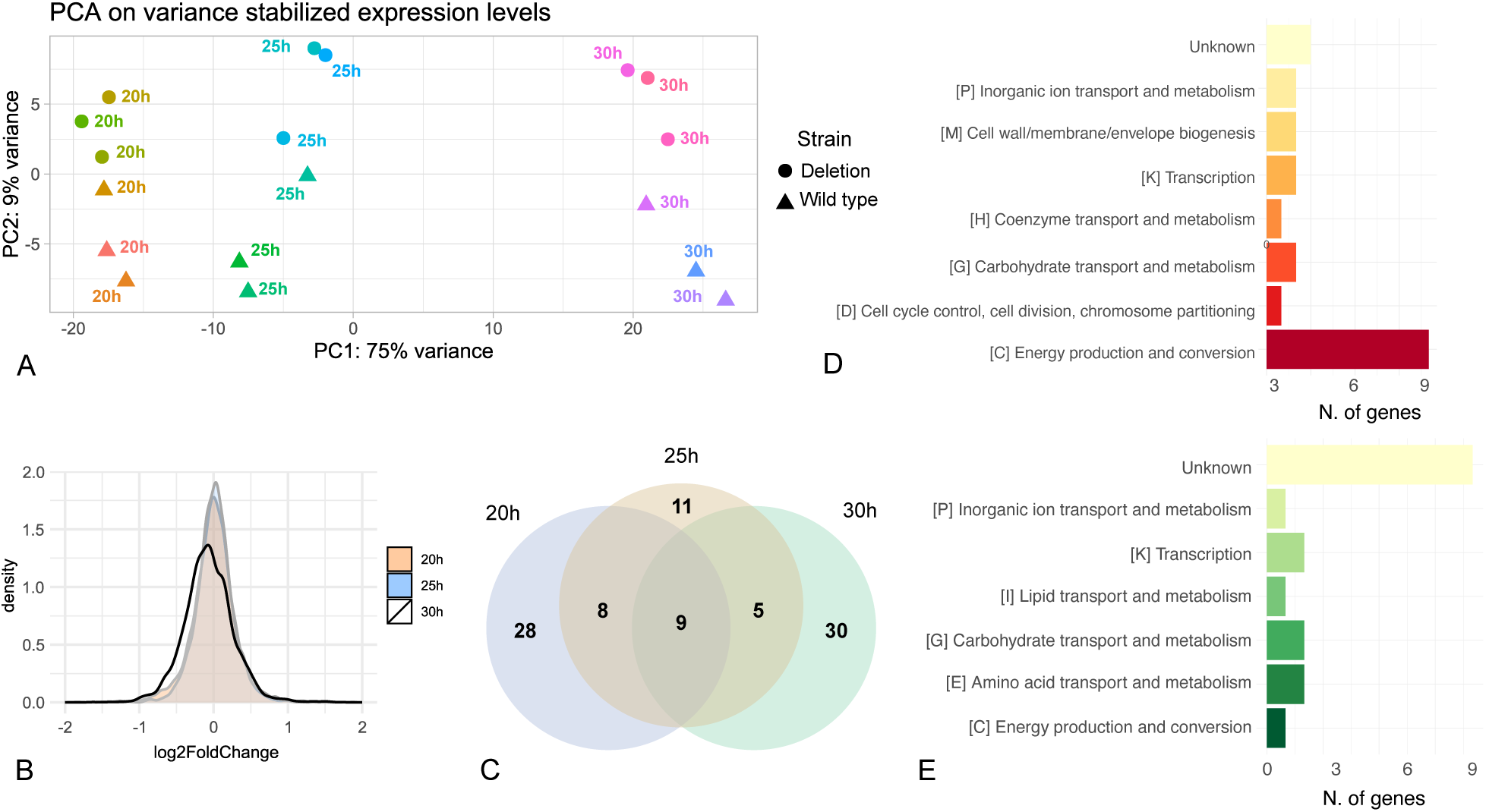
A) Principal component analysis on the scaled gene expression matrix of all samples. PC1: principal component 1. Principal components were calculated using plotPCA function from DESeq2 package implemented in R. B) Log2 fold change density chart of six samples per time point. Higher values indicate genes overexpressed in the deletion strain compared to the wild type. The figure cuts the LFC in the range [-2,2] since most of the data are centered in this interval. C) Venn diagram of DEGs at three time points, significant at padj*<*0.05 and *|l f c| >* 1 D) Functional categories of the upregulated genes in the mutant strains. E) Functional categories of the downrelgulated genes in the mutant strain

**TABLE 1.**
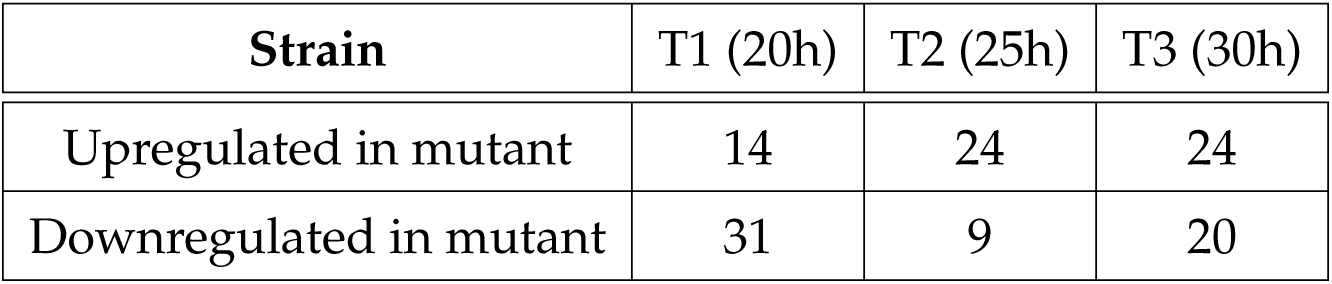
Number of up- and down-regulated genes in the mutant (*core* architecture) strain.

We then compared the set of DEGs of each time point and Figure 6C summarizes this analysis. Among the 91 distinct genes that passed our statistical test, 9 are shared by all three sets of DEGs. At 20h and 30h the majority of differentially expressed genes are proprietary to that specific time point, and the set of genes at T2 seems to be a transition state from a gene expression point of view (Figure **6**C). Interestingly, 5 out of the 9 genes shared by all samples, are involved in metabolism and, more specifically, in energy and carbohydrate metabolism (Table 2). Focusing on T3 (i.e. when, according to RNAseq and phenotypic data, the separation between *core* and *complete* strains is maximal), the involvement of metabolism is even clearer (Figure **6**D and E, Supplementary File S3 Table S3). More than half of the genes from both up-regulated and down-regulated categories belong to metabolic processes, with a predominance of energy metabolism among up-regulated ones and a more even distribution among metabolic functions in down-regulated ones. While the fraction of genes with unknown function is remarkable among down-regulated genes, these data indicate that one of the strategies with which the *complete* architecture achieves the buffering of noise of cell growth at the steady state could reside in the fine-tuned regulation of metabolism and, in particular, in the down-regulation of those involved in energy production. Indeed, these genes were found to be overexpressed following the removal of the *cciIR* module. The regulation of metabolism by means of QS has been widely described in microbes and it has been shown to play important roles in sustaining stable communities and shaping cooperation [30, 31]. Similarly to [32], our results suggest that the downregulation of key metabolic processes in a QS-dependent manner may represent a mechanism through which population-level homeostasis is achieved under crowded conditions. A possible scenario connecting gene expression and growth patterns observed here could envision the capability of the *complete* architecture to exert a tighter control over key nodes in energy metabolism (e.g. *acnB* aconitate hydratase 2 or D-isomer specific 2-hydroxyacid dehydrogenase only to cite the two top-ranking among upregulated DEGs) leading to a more uniform collective growth phenotype, possibly optimizing energy and resource utilization by individuals living in crowded environments. On the contrary, in the *core* harbouring cells, the absence of such metabolic breaks on the production of key intermediates might cause the loss of a coordinate response at increasingly higher cell densities. We emphasize that a mechanistic understanding of this specific process is lacking, and the data generated in this study will serve as a foundational contribution.

**TABLE 2.** The nine DEGs shared by the three sampling points of the RNAseq experiments whose expression changes significantly. *lcf* stands for log fold change.

| Locus Tag | lfc T1 | lfc T2 | lfc T3 | Annotation | Functional Category |
| --- | --- | --- | --- | --- | --- |
| K562_RS02985 | 2,49 | 3,39 | 3,46 | Chromate transporter | Inorganic ion transp./metab. |
| K562_RS26440 | 1,46 | 1,89 | 3,00 | *acnB* aconitate hydratase | Energy prod. and conv. |
| K562_RS30300 | 1,77 | 2,69 | 2,19 | Gluconolactonase | Carbohydrate transp./metab. |
| K562_RS30305 | 1,67 | 2,88 | 2,56 | 2-hydroxyacid dehydrog. | Energy prod. and conv. |
| K562_RS33000 | 1,08 | 2,62 | 1,73 | Tartrate dehydrogenase | Energy prod. and conv. |
| K562_RS03300 | -1,07 | -2,69 | -2,51 | Hypothetical protein | - |
| K562_RS03305 | -2,78 | -2,11 | -2,45 | Hypothetical protein | - |
| K562_RS03310 | -1,82 | -2,59 | -2,85 | VapC family toxin | - |
| K562_RS08800 | -1,06 | -1,94 | -1,59 | Putative lipoprotein | - |

## Discussion

QS is known to regulate hundreds of different traits in a given bacterial species [33]. QS regulatory circuits often display a complex organization that results in several feedback and feedforward loops [34, 35, 16, 36, 27]. Both the evolutionary history and the functional understanding of these complex structures are largely unknown [10]. Here we have shown that HGT is a major force driving the occurrence of multiple, integrated QS modules in the same genome [14]. In particular, the inclusion of additional regulatory systems (CciIR in the case of the *Burkholderia* genus) inside a genomic island represents an extraordinary example of the plasticity of the cellular transcriptional network and an exceptional occasion to study the effect of alternative regulatory architectures on the expression of specific phenotypic traits. Indeed, HGT and/or recombination events can lead to the appearance of novel regulatory modules (CciIR in our case) that must exhibit some degree of compatibility with pre-existing ones in order to be maintained over evolutionary time scales. [14]. To understand the functional implications of this increased structural complexity we modelled the additional control of *B. cenocepacia* K56-2 over the native CepIR system and simulated the functioning of this regulatory circuit. Guided by this theoretical framework, we experimentally showed that the simultaneous presence of the two QS modules can stabilize the end-point level of the population around a well-defined value. The lack of the non-native module, instead, introduces significant variability in cell density, starting from soon after the end of the exponential phase. Since it is known that quorum sensing heterogeneity is a feature associated with the low cell density state of bacterial populations [37], we confirmed the robustness of the *complete* circuit for initial cell density levels as low as OD=0.008. Interestingly, we didn’t notice any significant difference in the variability of *cepI* expression, at least at the population level. Whole transcriptome sequencing in the two strains, led to the identification of the cellular energy metabolism as a likely key factor in determining the two growth phenotypes observed here. Our analyses seem to suggest the capability of the *cepIR* regulatory module to fine tune key energetic pathways in the cell, whose deregulation may contribute to a non-cooperative growth behavior. Existing evidence shows that an imbalanced cellular metabolism could lead to noisy cell density patterns (and *viceversa*) and this, in turn, may be caused by a plethora of different mechanisms. Metabolic upregulation, for example, may generate end products that could interfere with overall gene expression and cellular activity thus introducing feedback mechanisms that introduce fluctuations that contribute to cell density noise [38, 39]. Moreover, feedback regulation of glycolitic enzymes can lead to oscillations in metabolic activity that influence growth rates and cell density stability [40]. Furthermore, metabolic dysregulation could impact the biosynthesis of QS autoinducers, thereby disrupting coordinated and homogeneous population-level growth phenotypes.

Through computational analysis, mathematical modeling, in vitro experiments and whole genome and transcriptome sequencing we find and establish a connection between quorum sensing, metabolism and fine-tuning of bacterial population density in *Burkholderia*, yet some aspects of the more intimate relationship of these three players remain obscure. For example, although we didn’t observe any population-level heterogeneity of *cepI* levels, it remains to be addressed whether the expression of this AI-synthase occurs stochastically at the single cell resolution (as observed in other organisms in [26, 41]) and/or whether ii) this is somehow affected by the two different QS regulations configurations. It would also be intriguing to quantify the concentration of key intracellular energetic metabolites and assess whether the observed differences in specific metabolic genes propagate throughout the entire metabolic network, and if so, to what extent. Finally, since it has been shown that CepR and CciR are capable of jointly tuning the expression of many virulence factors [42], we acknowledge the relevance in pointing future work toward the characterization of key pathogenic phenotypes in relation to the two distinct architectures investigated in this study, and whether or not they represent actionable target for new therapeutic strategies. Further work is necessary to address these specific questions.

## Methods

### Retrieval of genomic islands

We retrieved genomic islands information from the IslandViewer4 database [43], maintained by the Brinkman lab at Simon Fraser University, Canada. The pre-computed genomic island coordinates for all sequenced genomes until March 2022 were downloaded from IslandViewer at *pathogenomics*.*s f u*.*ca*/*islandviewer*/ on July 29 2023. After much research, we have chosen this framework because of its consistency in providing updated GI predictions over several years and because it offers information based off three diverse prediction methods. Briefly, SIGI-HMM [44] identifies alien genes using a generative model on codon usage bias, bypassing the highly expressed genes. IslandPath-DIMOB [45] couples dinucleotide bias and mobility gene presence in a window of at least eight genes as a sensitive way to detect GI regions, whereas IslandPick [46] finds probable islands and non-island regions through a flexible comparative genomic approach. We downloaded the data set that is the union of SIGI-HMM, IslandPath-DIMOB and IslandPick to try capturing as much variability as possible. We refer to this as the IslandViewer4 data set, that is a table of genomic coordinates (see **Supplementary File S1**).

### Retrieval of genome sequences and annotations

The IslandViewer4 data set contains chromosome names, but it does not tie them to the assembly accession they belong to. We therefore first downloaded the assembly structure reports files sourced by RefSeq from *https*: //www.ncbi.nlm.nih.gov/assembly by searching *^t^ Bacteria^t^* and applying the filters "Complete genome" and "Chromosome", to be consistent with [43]. On Sept 12*^th^* 2023 this search produced 40255 results, which we downloaded and copied to our local server. These files were parsed and a conversion table associating assembly accession and chromosome names was extracted. Crossing the conversion table with the actual chromosome names in the IslandViewer4 data set allowed us to retrieve a unique list of 17868 assembly accession numbers used in the prediction of the genomic islands by the above mentioned third party. We used the NCBI datasets standalone application to download the sequences and annotations of the bacterial genomes present in the unique list of accessions derived from the IslandViewer4 data set. The list of accessions is available in **Supplementary File S2**.

### Identification of LuxIR-like sequences and data validation

After the pre-identification with HMMER we further validated our data having all analyses active on the InterProScan software, and filtering for an E value of at most 0.001. We labelled as P (passed), the LuxI-like proteins predicted as IPR001690 (Autoinducer synthase), IPR016181 (Acyl-CoA acetyltransferase) or IPR018311 Autoinducer synthase CS). We also labelled as P the LuxR-like representatives predicted to have both the autoinducer-binding and the helix-turn-helix domain, formally IPR005143+IPR000792, or IPR036693+IPR016032 or IPR036693+IPR036388. We did not discard the entries that did not pass the InterProScan validation, but we labelled them as F (failed). The data table is available in Dataset S1. Using these filters, we obtained an *F*/*P* ratio of 0.40. Throughout the manuscript we refer to LuxIR-like as only the proteins that passed the InterProScan validation.

### ANI calculation

The relatedness of *Burkholderia* genomes was measured using Average Nucleotide Identity (ANI) as implemented in FastANI [47].

### Bacterial strains, media and culturing conditions

The bacterial strains and plasmids used in this work are reported in Table S4. Unless stated otherwise bacteria were grown under aerobic condition at 37 °C in Luria-Bertani (LB) agar or broth. Antibiotic concentration used were 40 *µ*g/ml kanamycin, 50 *µ*g/ml trimethoprim, 35 *µ*g/ml chloramphenicol and 15 *µ*g/ml tetracycline for *E. coli*, 100 *µ*g/ml trimethoprim, 200 *µ*g/ml chloramphenicol, 200 *µ*g/ml ampicillin and 150 *µ*g/ml tetracycline for *B. cenocepacia* K56-2. All antibiotics were purchased from Merk Life Science S.r.l..

### Construction of deletion strains

Deletion strains were constructed as described in [48, 49]. Briefly, two DNA regions flanking the CciIR system genes were amplified from the *B. cenocepacia* K56-2 genome using the primer pairs cciI_L_for-cciI_L_rev and cciR_R_for-cciR_R_rev (**Table S5**). PCR amplifications were performed using Phusion^TM^ High-Fidelity DNA Polymerase (Thermo Fisher Scientific) with specific amplification conditions for each primer pair. The obtained PCR products were digested with XbaI-BamHI and BamHI-KpnI (Thermo Fisher Scientific) respectively and ligated (T4 DNA Ligase, Thermo Fisher Scientific) into the pGPI-SceI-XCm suicide vector (containing a unique restriction site for the endonuclease I-SceI and the *xylE* reporter gene, **Table S**4) [48] digested with XbaI-KpnI. The obtained plasmids were introduced by transformation in *E. coli* SY327 and then mobilized by conjugation to *B. cenocepacia* K56-2. Exconjugants were selected in the presence of trimethoprim, chloramphenicol and ampicillin at the concentrations reported above for each strain. Then, a second plasmid, pDAI-SceI-SacB (encoding the I-SceI endonuclease) [48] was introduced by conjugation, producing site specific double-strand breaks at the I-SceI recognition site. Resulting exconjugants were tetracycline resistant (due to the presence of pDAI-SceI-SacB) and trimethoprim and chloramphenicol susceptible (indicating the loss of the integrated mutagenic plasmid) and were selected using tetracycline and ampicillin at the concentrations reported above for each strain. The loss of the integrated plasmid was confirmed by plating in presence of trimethoprim and chloramphenicol. Finally, the plasmid pDAI-SceI-SacB was cured, and cured mutants were selected on LB plates without salt and supplemented with 5% (*wt*/*vol*) sucrose (Oxoid S.p.A.) and then screening the resulting colonies for loss of tetracycline resistance. The desired gene deletions were first confirmed by PCR amplification using primer pair E_cciI_for-E_cciR_rev (Table S5) and then by genome sequencing.

### Genome sequencing

Genomic DNA of *B. cenocepacia* K56-2, K56-2 Δ *cciIR*, K56-2 *pcepI-lux* and K56-2 Δ*cciIR* pcepI-lux was extracted using the DNeasy UltraClean Microbial Kit (Qiagen S.r.l). Whole genome shotgun sequencing (2 x 150 bp) was performed by BMR genomics S.r.l.. The sequencing reads were submitted to the NCBI on Oct-30-2023, and are available under the SRA experiment identifier PRJNA1033778.

### Measurement of the activity of pcepI-lux reporters

To determine the levels of expression from the promoter regions of *cepI* and growth dynamics in the *B. cenocepacia* K56-2 p*cepI*-lux and K56-2 Δ*cciIR* pcepI-lux strains, overnight culture of each strain were diluted until an OD600 comprised between 0.008 and 0.02 (depending on the experiment) and 200 *µ*l of this dilution were aliquoted in triplicate in a white assay 96 wells microplate with clear bottom (VWR International S.r.l.) and incubated at 37°C in an Infinite 200 PRO Tecan plate reader. Both OD600 and luminescence were measured every hour for 48h. Luminescence was expressed in relative light units per optical density of the culture (RLU/OD600).

### Growth curve experiments for strains AU1054 and J2315

For growth curves of *B. cenocepacia* J2315 and AU1054 strains, overnight cultures of each strain were diluted until an OD600 between 0.008 and 0.02 and 200 *µ*l of this dilution were aliquoted in triplicate in a 96 wells microplate (Sarsted AG and Co.) and incubated at 37°C in an Infinite 200 PRO Tecan plate reader. OD600 was measured every hour for 48h.

### Mathematical modeling and parameters fitting

The mathematical model (both deterministic and stochastic) of the QS regulation in *B. cenocepacia* K56-2 is reported in detail in **Supplementary File S3**. The deterministic model described therein was implemented using MATLAB^®^ 2022b. The *ode45* solver was used to solve the set of differential equations accounting for the dynamics of the system. To estimate the parameters of the model from experimental data we used a stochastic curve-fitting in-house MATLAB software. The algorithm is based on the paper by Cardoso et al. 1996 [50] and consists in the combination of the non-linear simplex and the simulated annealing approach to minimize the squared deviation function. The set of fitted parameters are reported in Table S1 and S2, together with their confidence intervals (CI). 95% CI of the fitted parameters were computed by performing 100 bootstrap resamplings of the original data matrices and estimating, for each of them, the best parameters set. Values presented are averages of these 100 resemplings. Finally, the stochastic simulations were performed implementing the Gillespie algorithm [24] in C. The codes used to perform the simulations reported in this work are available at *DOI*: 10.5281/*zenodo*.12796961.

### Sampling for RNA sequencing experiment

For RNA-seq, overnight cultures of *B. cenocepacia* K56-2 and K56-2 Δ*cciIR* were diluted until an OD 600 of 0.01 and 200 *µ*l of this dilution were aliquoted in quadruple in a 96 wells microplate (Sarsted AG & amp; Co.) and incubated at 37 °C in an Infinite 200 PRO Tecan plate reader (Tecan Group Ltd.). OD 600 was measured every hour. After 20h, 25h and 30h, cells from each of 4 replicates of the same strain, were recovered and pooled, and 500 *µ*l of this pool were treated with the RNA protect bacteria reagent (Qiagen S.r.l.) and conserved at -80 °C. Each experiment was repeated three times.

### RNA extraction and sequencing

Total RNA was extracted using a RNeasy Mini Kit (QIAGEN S.r.l.) following manufacturer’s instructions, extending the incubation with the DNase enzyme (RNase-Free DNase Set, QIAGEN S.r.l.) to 1 hour. RNA concentration and quality were assessed using a QUBIT RNA assay kit and a Qubit 4 Fluorometer (both from Invitrogen-Thermo Fisher Scientific Inc.) and an Agilent RNA 6000 Nano kit and an Agilent 2100 Bioanalyzer (both from Agilent Technologies Italia S.p.A.). RNA sequencing (2 x 150 bp) was performed by BMR genomics S.r.l. on Illumina platform (Illumina, Inc.). The sequencing reads were submitted to the NCBI and are available under the SRA experiment identifier PRJNA1033778.

### Sequencing reads filtering and validation

Quality assessment and filtering using fastp 0.23.2 (parameters: –cut_tail –average_qual 20 –trim_poly_g –detect_adapter_for_pe –length_required 55 –overrepresentation_analysis –overrepresentation_sampling 1000) yielded most of the initial reads, with very little trimming and filtering needed. The overrepresentation flag enabled the sampling of one every 1000 sequencing reads, which were later confirmed to be mostly ribosomal RNAs.

### Sequencing reads diagnostics

RefSeq (GCF_014357995.1) *Burkholderia cenocepacia* K56-2 reference genome, GFF3 annotation and transcripts files were accessed on 18th March 2024 via NCBI. Polished reads were mapped to the reference genome using bowtie2 2.4.4 in sensitive mode and enabling the end to end setting (entire read must align, parameters: –end-to-end –sensitive). Inspection of the SAM files showed *>*98% of reads mapping to the reference genome sequence with an equal balance between read pairs flagged as 99/147 (properly paired, R1 aligns to the forward strand) and 83/163 (properly paired, R2 aligns to the forward strand), consistent with the equal strandedness of the reference genome and our inward stranded reverse library preparation protocol. Half or more of the read pairs mapped to non-coding RNAs (mostly rRNA), and this was also confirmed by a manual check of the overrepresented reads after filtering with fastp. A median of 99.03% reads aligned well to the reference genome, confirming no major flaws in sequencing or in the experimental procedure.

### Transcripts abundance estimation

An index file for quantification was built from the RefSeq transcript file using salmon version 1.10.0 using k-mer=21 and with decoy awareness enabled (parameters-k and-d). Transcripts abundance estimation was performed with salmon 1.10.0 in selective alignment mode using the inward stranded reverse parameter “ISR”, consistent with our library preparation layout (parameters:-l ISR –seqBias –gcBias –posBias –incompatPrior 1 –minScoreFraction 0.8 –numGibbsSamples 10). We enabled these command line options to fine-tune the quantification process and account for various sequence-specific biases and improve accuracy in estimating transcript abundances. A total of 292,463,702 raw read counts were estimated across 18 samples and 7,008 transcripts. The final number of genes effectively utilized (row sum over all samples below 20) was 6,974, with locus tag prefixes “K562_RS”. An average of 44.16% (s.d 10.48) of the total reads were attributed to mRNA, which is expected since the samples were dominated by non-coding RNAs, which the NCBI transcripts file omits. The codes used for the analysis of the RNAseq data are available at *DOI*:10.5281/zenodo.12796961.

## Acknowledgements

C.R. expresses his gratitude to Justin Cook at Simon Fraser University in relation to key additional details about IslandViewer4.

## DATA AVAILABILITY STATEMENT

All data and code available at 10.5281/zenodo.12796961.

## FUNDING

F.D.P. acknowledges the funding from Project ‘Mathematical modelling for a sustainable circular economy in ecosystems’ (grant n. P2022PSMT7) funded by EU in NextGenerationEU plan through the Italian ‘Bando Prin 2022-D.D. 1409 del 14-09-2022’ by MUR. M.F. acknowledges the funding from Project "EXPLORE-EXploiting pathogens PLOidy to fight drug REsistance: towards a precision medicine approach" funded by EU in NextGenerationEU plan through the Italian ‘Bando Prin PNRR 2022-D.D. 1409 del 14-09-2022’ by MUR.

## CONFLICTS OF INTEREST

The authors declare no conflict of interest.

